# Disentangling maternal pregnancy and fetal-specific plasma signatures through integrated proteomics and site-specific N-glycoproteomics

**DOI:** 10.64898/2026.10.05.756652

**Authors:** Shelley Jager, Sofia Kalaidopoulou Nteak, Grigory Reznikov, Francine Rodrigues Ianiski, Remco Visser, Pavel Sinitcyn, Gestur Vidarsson, Albert J.R. Heck

## Abstract

During pregnancy, mother and fetus co-exist albeit as two physiologically distinct individuals, connected through the placenta and the umbilical cord. This connection allows for the exchange of vital substances, while individual blood homeostasis remains intact. We hypothesized that comparing maternal and cord plasma (N-glyco)proteomes would provide insights into the biological communication and unique physiological state of each individual. As comparing only matched maternal and umbilical cord blood plasma misses a crucial identifier in determining whether observed changes are related to infancy, pregnancy or both, we designed a triad study by adding a non-pregnant women control group. This allowed us to distinguish pregnancy-specific proteins and their potentially altered N-glycosylation from those in infancy. We found that the fetus and mother have distinct proteomes, N-glycoproteoforms and alternative complement regulation. Moreover, we observed that pregnancy in women led to specific N-glycosylation changes on a select set of primarily HDL-associated and acute-phase proteins. Lastly, our data provide evidence for an early fetal development of the humoral immune system as especially fetal IgM displayed distinctive N-glycosylation signatures.

## Introduction

In pregnancy, mother and fetus are in a unique physiological state where two genetically distinct individuals coexist, while remaining connected through the placenta and the umbilical cord. Throughout gestation, the mother and fetus exchange nutrients, hormones, metabolites and immunological signals to support the fetal development while preserving maternal homeostasis^1,2^. This process reshapes the circulating plasma proteome for both fetus and mother^3–6^. Characterizing these changes is essential for understanding normal pregnancy and provides the foundation for identifying deviations associated with maternal and neonatal diseases.

These physiological adaptations are reflected in the protein composition of blood plasma, which provides an accessible window into systemic physiology^7^. As the primary transport medium for circulating proteins, plasma integrates changes in immune regulation, coagulation, lipid metabolism, and intercellular signaling that accompany both health and disease^8^. In pregnancy, the maternal plasma proteome undergoes dynamic remodeling throughout gestation, while the protein composition of the umbilical cord plasma changes during fetal development and reflects the physiological state of the newborn at birth^3–6^. In addition, maternal and fetal blood are connected through the placenta^1^. Consequently, analyzing the maternal and cord plasma proteomes can provide insights into the biological communication between mother and fetus.

Changes in plasma proteome physiology are not solely determined by protein abundance. Especially for plasma proteins, N-glycosylation plays an important role in their function, stability and molecular interactions^9–12^. Furthermore, the establishment of healthy maternal-fetal crosstalk is mediated by N-glycosylation^13,14^. Pregnancy-associated alterations in overall N-glycosylation have been reported previously, based on released N-glycan analyses^14^. Additionally, N-glycan profiling for several individual plasma proteins has been performed, including immunoglobulins^15,16^, alpha-1-antitrypsin^17^, and alpha-1-acid N-glycoprotein 1^18^, with the latter two exhibiting increased abundances of tri- and tetra-antennary N-glycans related to pregnancy. In addition, released-N-glycan analyses revealed that maternal and umbilical cord plasma display overall different N-glycan-signatures with umbilical cord plasma relatively enriched in bi-antennary N-glycans, whereas maternal plasma contained a higher proportion of branched and highly sialylated N-glycans^19^. However, released-N-glycan analyses cannot identify the proteins or N-glycosylation sites responsible for these differences, leaving it unclear whether pregnancy-associated N-glycan remodeling reflects changes in a limited number of abundant N-glycoproteins or whether these originate from widespread alterations across the plasma N-glycoproteome. Moreover, comparisons restricted to matched maternal and umbilical cord blood plasma cannot distinguish whether observed molecular differences arise from fetal physiology or from the profound remodeling of the maternal plasma proteome and N-glycoproteome during pregnancy. Including healthy non-pregnant women as a physiological reference base provides the necessary framework to disentangle pregnancy-associated adaptations from fetal-specific molecular traits.

Therefore, here we performed proteomics and N-glycoproteomics on a cohort of matched umbilical cord plasma and maternal plasma as well as plasma from a non-pregnant women control group (**Figure 1**). This triad design enabled us to assign molecular traits to normal female physiology, pregnancy-associated adaptation, and fetal biology, while simultaneously identifying shared and distinct protein abundance and protein-specific N-glycosylation patterns.

**Figure 1:**
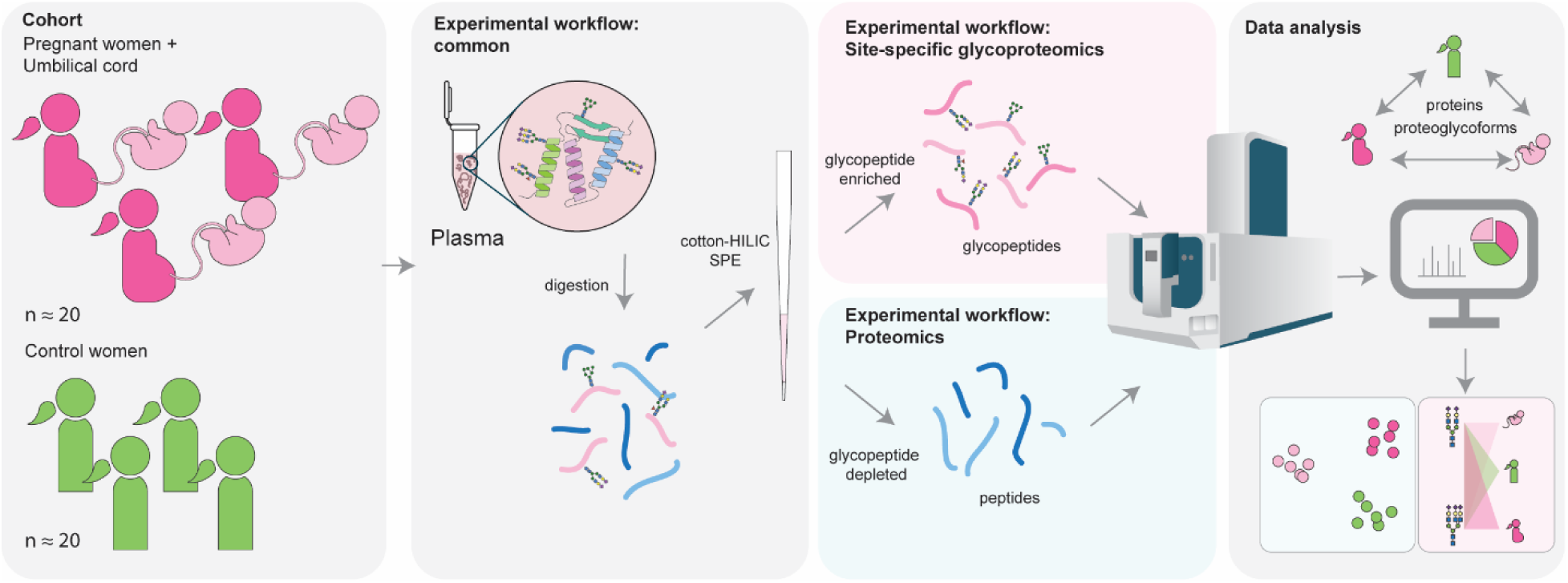
The triad study design and experimental approaches. The panel on the left portrays the study cohort. The cohort encompassed ∼20 matched mother-umbilical cord pairs. To complement this cohort, 20 plasma samples obtained from healthy non-pregnant women were added as a control group. The next panels depict the experimental workflow for the N-glycoproteomics and proteomics samples: plasma was digested into peptides and subsequently enriched using cotton-HILIC SPE. The resulting enriched N-glycopeptides (pink panel) were analyzed using a N-glycoproteomics workflow, while the remaining N-glycopeptide depleted pool was used for standard proteomics (blue panel). Pink and blue colors are used throughout the manuscript to describe N-glycoproteomics data or proteomics data, respectively. Peptides were analyzed by mass spectrometry and data from proteomics and N-glycoproteomics were analyzed using dedicated computational workflows (right panel), to derive proteomics and N-glycoproteomics signatures defining maternal or fetal physiology.

## Results and discussion

### Proteomics analysis revealed distinct features for fetus as well as pregnant women

At first, we examined the proteomes of each individual using label-free quantitative proteomics, following procedures and protocols described previously^20^. A more detailed description of the cohort is provided in Supplementary Table 1 and 2. Using the label-free DIA-based proteomics approach, we were able to identify and confidently quantify around 250 different proteins in each plasma sample.

To identify specific protein signatures that define each physiologically distinct group, we first explored the quantitative proteomics data by hierarchical clustering (**Figure 2A**). This clustering revealed a clear and strong separation between the three groups. The fetal group stood out the most, displaying a very distinct proteome with a large group of proteins that are either highly enriched (including the fetal protein marker AFP (alpha-fetoprotein) and various collagen-alpha chains (COL1A, COL2A1, COL5A1, COL6A1 and COL6A3)) or highly decreased (including HP (haptoglobin), IGHA1 and IGHA2 (immunoglobulin heavy chains A1 and A2) (clusters I and II, respectively; **Supplemental Table 3**)). The proteins in cluster I and II displayed comparable abundances between the pregnant and control women, which further showed that the alternative regulation was specific to the fetal physiology. GO-term enrichment (**Figure 2B and 2C**) revealed that both upregulated and downregulated proteins could be classified into the terms humoral immune response and complement activation pathway, suggesting substantial alternative regulation in these two pathways for infants. In addition, the downregulated proteins (cluster II) in the umbilical cord plasma were found to be involved in the acute-phase response.

**Figure 2:**
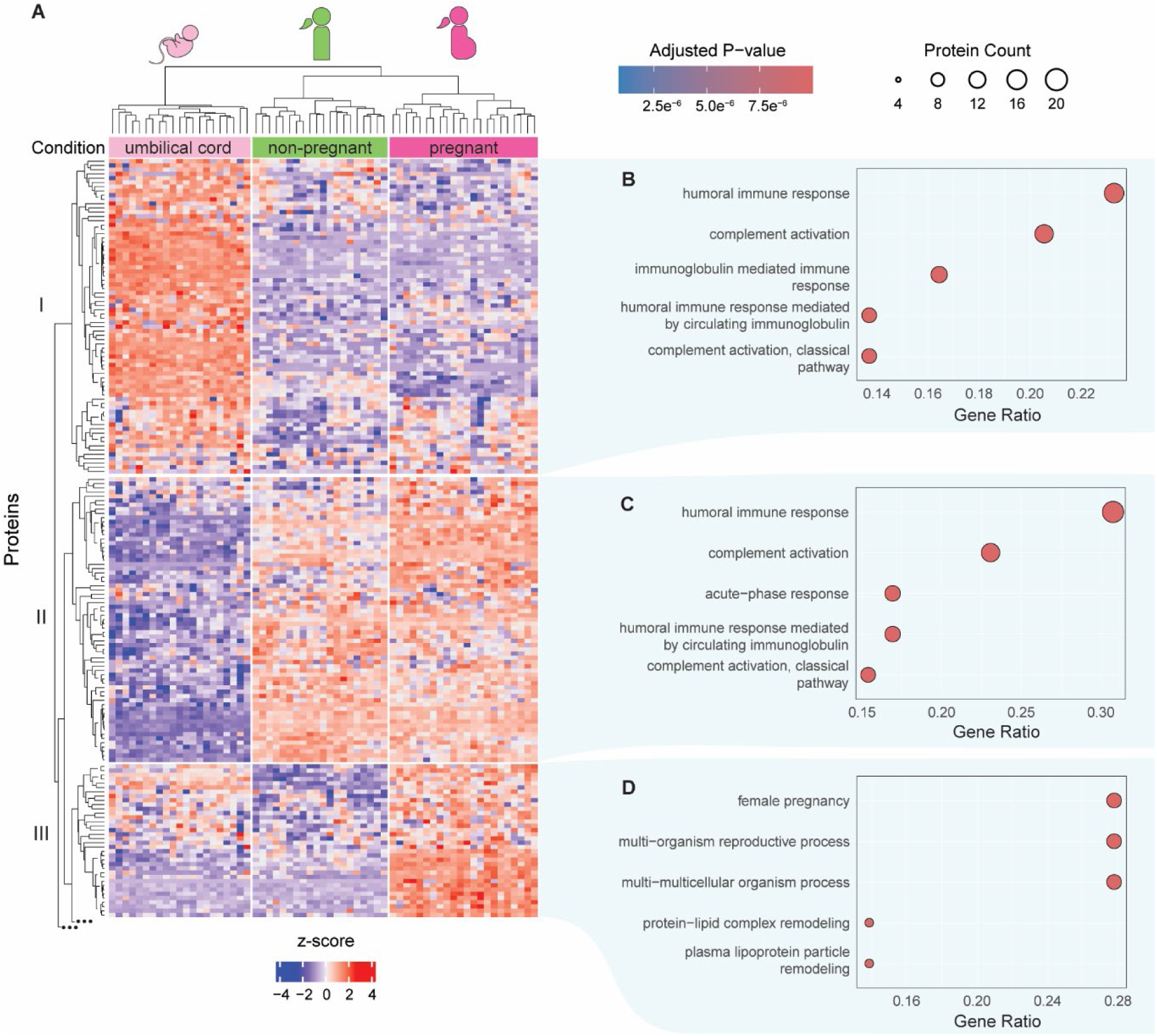
Proteomics analysis reveals distinct protein clusters, particularly for fetal plasma. **A)** Hierarchical clustering provides clear clusters distinguishing the three groups (non-pregnant women, pregnant women and fetus (umbilical cord). The three protein clusters shown in this heatmap I (74 proteins), II (67 proteins), and III (36 proteins) represent the largest clusters. They represent the upregulated (I) and downregulated (II) proteins in the fetus, as well as the pregnancy-related proteins (III) that were found to be more abundant exclusively in pregnant women. Z-scores were calculated separately for each protein across all samples using the log2-transformed LFQ intensities. In total, unsupervised clustering revealed 6 clusters; the remaining clusters are removed for simplicity (represented with the three dots). The complete heatmap of all proteins is provided in Supplemental Figure 1, and all proteomics data is available through Supplemental Table 3. GO enrichment analysis of **B)** the cluster I proteins, **C)** the cluster II proteins, and **D)** the cluster III proteins. The color of the dots represents the adjusted p-value and the size the protein count.

Another clear cluster was cluster III. Cluster III is specific to pregnant women and was found, as expected, enriched in pregnancy-related proteins (PZP, PAPPA, several PSG proteins, LNPEP, ADAM12, and SVEP1) (**Figure 2D**; **Supplemental Table 3**). Furthermore, there is a group of proteins that has decreased abundance in both the pregnant women and the fetal group (Cluster V, **Supplemental Figure 1B)**. These proteins were found to be related to zymogen activation, cell-cell adhesion and fibrinolysis, suggesting modulation in these pathways for both mother and infant.

Next, we zoomed in on the proteins involved in complement activation. This process was highly enriched in the GO-term analysis of both clusters I and II, suggesting alternative regulation in the neonate. However, looking at the complement proteins revealed that pregnant women also exhibit different protein abundances compared to the control women **(Supplemental Figure 2)**. Complement activation is initiated when pathogen-associated molecular patterns are recognized by so-called pattern recognition molecules (PRMs)^21^. A cascade of consecutive activation through the complement proteins leads to the formation of the membrane attack complex (MAC), which can open pores in pathogenic cells. There are several other cofactors and soluble proteins that interact with the complement system, both as positive and negative regulators. A simplified overview of the complement activation cascade and the average protein abundances as observed in each group is illustrated in **Supplemental Figure 2B-D**.

In the fetus, we observed that proteins involved in PRM-complexes of both the classical and the lectin pathways are relatively more abundant compared to both adult women groups (**Supplemental Figure 2B-D**). The same was found for ficolin-3 (FCN3), which activates the lectin pathway. In addition, complement factor C2 and both C4A and C4B are also relatively more abundant in umbilical cord plasma compared to both adult women groups, while the C4 inhibitors (C4BPA and C4BPB) displayed substantially decreased abundance in umbilical cord plasma. In summary, the initiators of the complement pathway are more abundant in the umbilical cord plasma, while the inhibitors are less abundant. Conversely, proteins involved in the MAC complex are less abundant in umbilical cord plasma.

In the pregnant women’s plasma, the initiators in the classical and lectin pathway were slightly more abundant compared to the non-pregnant women, but so were the C4 inhibitors. Interestingly, we also observed a slight increase in CRP, reflecting a slightly elevated inflammatory state. Another very pronounced feature in pregnant women is the upregulation in complement factors involved in the MAC complex (C5, C6, C8, and C9, **Supplemental Figure 2C**), which we do not observe in umbilical cord plasma or non-pregnant women.

In pregnancy, the maternal immune system must defend against pathogens but must simultaneously tolerate the fetus^22^. In agreement with our data, previous studies have reported increases in protein concentrations of the negative complement regulator factor H throughout pregnancy^23,24^. Furthermore, normal pregnancy has been associated with the activation of the classical and lectin pathways and increased formation of the terminal complement complex^25^. Complement activation is also relevant to placental development, where moderate activation and a slightly inflammatory state, reflected by increased CRP, may support normal development. Excessive activation, on the other hand, has been associated with adverse pregnancy outcomes. Our findings also indicate remodeling of the complement pathway, in particular, we noticed increased concentration of several MAC-components in pregnant women.

Due to the increased complement activation and inflammatory state of the mother, the fetus needs a protective mechanism against maternal complement attack^26,27^. In early pregnancy, sialic acids on trophoblasts have been proposed to protect the embryo from such threats, while loss of sialylation can promote activation of the maternal alternative pathway in mice^27^. It is not known whether this is relevant in late pregnancy, the timepoint at which the here used samples were acquired.

Furthermore, the fetal complement profile was distinct from that of both pregnant and non-pregnant women, and was characterized by increased abundance of proteins involved in initiating the classical and lectin pathways and the decrease of many complement inhibitors. This may reflect the developing and naive fetal immune system, which might require different and more sensitive regulation to respond to threats.

### The plasma N-glycoproteome is heavily affected by pregnancy

In addition to the proteomics data, we recorded N-glycoproteomics data on the same samples. Cumulatively, we detected 117 N-glycoproteins, harboring 228 distinct N-glycosites, and around 2000 unique N-glycopeptides. Of these, on 66 proteins and 104 N-glycosites we identified three or more different N-glycan compositions, which we used for the analyses. Depending on the N-glycosite, we were able to identify up to 40 different N-glycans per site.

To grasp the heterogeneity of these site-specific N-glycosylation features, we first asked which N-glycosites carry similar N-glycosylation within a single group. Instead of exploring the N-glycosylation for each N-glycan separately, we compared the full N-glycan distribution on a single site between donors (which we will refer to as the N-glycosylation profile), using cosine distances (e.g. site 1 versus site 2, both in pregnant women). If the cosine distance is close to the maximum of 1, it means two individuals have a very different N-glycosylation profile; while if the cosine distance is close to 0, it means two individuals have a highly similar N-glycosylation profile. We summarized these findings in a clustering dendrogram for each group, as shown in **Supplemental Figure 3**. This revealed that many N-glycosylation sites actually have the same N-glycosylation profile in a single physiological group. Interestingly, the N-glycosites that were grouped together in the non-pregnant women, and thus had a similar N-glycosylation profile, were not always the same N-glycosites that grouped together in the umbilical cord plasma group or in the pregnant women group (see **Supplemental Figure 3**).

To dive deeper into this, we first performed a side-by-side comparison of the clusters of the control women and the fetus, to see which N-glycosites no longer clustered with each other (**Supplemental Figure 4**). We only examined the N-glycosites that were detected in both groups and belonged to the major clusters in either group (with 3 or more N-glycosites). Interestingly, while these N-glycosites formed the 4 large clusters in the control group, they only formed 2 major ones in the fetal plasma. This decrease in the number of clusters already indicated that more N-glycosites in umbilical cord plasma of different donors carry similar N-glycan profiles to one another than in either adult group. The largest group in the non-pregnant women consisted of 40 N-glycosites, and the N-glycosylation profile was dominated by N4H5S2 (**Supplemental Figure 4**, green panel), which is well-known to be the most frequently detected and abundant N-glycan in human plasma. These same N-glycosites also belong to the largest cluster in fetus, and the average N-glycan profile is also the same. The next largest cluster in non-pregnant women (cluster 2) consisted of 5 N-glycosites. The N-glycosylation profile on the sites belonging to cluster 2 was dominated by less sialylated di-antennary N-glycans (mainly N4H5S1). The N-glycosylation profile of cluster 3 in non-pregnant women was dominated by two N-glycans: N4H5S2 and N5H6S3 (a tri-antennary N-glycan species). Interestingly, clusters 2 and 3 did not exist in the fetus. Instead, several sites actually clustered together with sites in cluster 1, representing that their N-glycosylation profile shifted to the N4H5S2 dominant profile. In the final main cluster (cluster 4), the N-glycosylation was mostly dominated by oligomannose-type N-glycans (N2H5-N2H8), both in the control women and in the umbilical cord plasma. The N-glycosites in this cluster remained clustered together in the fetus. In summary, comparing the N-glycosylation profile clustering of non-pregnant women to that in the fetus revealed that fewer clusters in the fetal plasma. Thus, in the fetus, N-glycosylation is more similar across different N-glycosylation sites which are mainly occupied by N4H5S2 (a di-antennary, fully sialylated, complex-type N-glycan).

We performed a similar side-by-side analysis of the N-glycosites clusters of non-pregnant women and pregnant women (**Figure 3A**). This revealed again that several N-glycosites that had similar N-glycosylation profiles in non-pregnant women, no longer clustered together in pregnant women. While in the previous comparison (non-pregnant women vs. fetus), N-glycosites were more di-antennary, we observed a different change when comparing non-pregnant women to pregnant women: several N-glycosites clustered with the tri-antennary dominant cluster (cluster 3) instead. In other words, these N-glycosites change from a dominant di-antennary N-glycan profile to a dominant tri-antennary N-glycan profile. Linking this to the N-glycoproteins, these sites include all identified sites of kininogen 1 (KNG1), and one site of alpha-1-acid N-glycoprotein 1 (ORM1), one on alpha-1 antitrypsin (SERPINA1), one on plasma protease C1 inhibitor (SERPING1), and one on vitronectin (VTN). All these proteins are mainly produced in hepatocytes^9^. The co-occurrence of these shifts therefore points to a change in the N-glycosylation of hepatocyte-derived plasma proteins during pregnancy.

**Figure 3:**
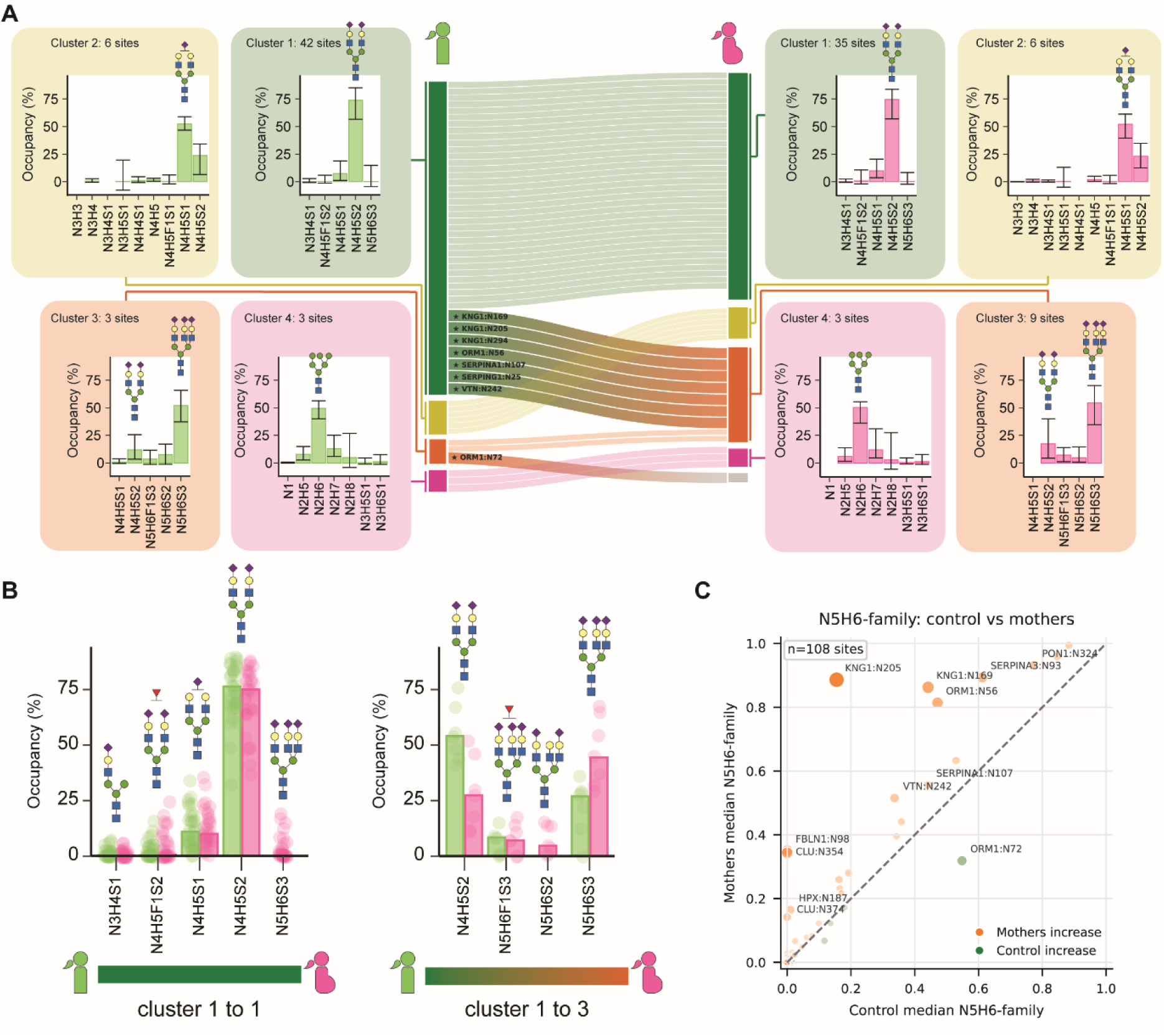
Pregnancy leads to a relative increase in the abundance of tri-antennary N-glycans on a specific subset of N-glycosites and N-glycoproteins. A) Sankey diagram displaying how N-glycosites cluster based on N-glycosylation similarity. Each line represents a single N-glycosite. Four major clusters with multiple N-glycosites could be categorized. Sites belonging to a single cluster in non-pregnant women were observed to move to another cluster in pregnant women. The N-glycosites that shift are annotated with their gene names and N-glycosite amino acid number. On each site, the average N-glycan distribution across all N-glycosites in clusters 1 to 4 is shown. B) Side-by-side N-glycan distributions of the N-glycosites that retain in cluster 1 (left) and those that move from cluster 1 to 3 in pregnancy (right). In A and B, error bars represent the 25^th^ and 75^th^ percentiles, and major N-glycoforms are annotated using cartoons. C) Correlation plot of the median relative abundance of N-glycoforms belonging to the N5HC family (N5HCF_x_S_y_) in pregnant (y-axis) and non-pregnant women (x-axis), to represent increased tri-antennary N-glycans at several N-glycosites. Point size represents magnitude of the change, and dots furthest from the diagonal are annotated with gene name and N-glycosite.

Interestingly, further examination of these seven sites revealed that they already had a higher abundance of tri-antennary N-glycosylation in the non-pregnant group compared to the sites that remained in cluster 1 (**Figure 3B**). In the non-pregnant women, the abundance of these tri-antennary N-glycans is generally lower than their di-antennary counterparts. In pregnancy, however, the tri-antennary species becomes the dominant N-glycan specifically on these shifting sites. The observed change is selective rather than global, as most N-glycosites on other liver-derived N-glycoproteins still cluster together in cluster 1 and present the same average N-glycosylation profile as in non-pregnant women.

To further confirm that the shift towards tri-antennary N-glycans was selective to only a few N-glycosites, we compared the ratio of tri-antennary species in pregnancy to those in the control group of all N-glycosites (**Figure 3C**). Because the data obtained here is purely compositional and not structural, we inferred the antennarity based on the composition. Therefore, we assumed that all N-glycans with 5 N-acetyl hexosamines and 6 hexoses (N5H6) are tri-antennary. This analysis confirmed again that the previously mentioned shifting N-glycosites (e.g. KNG1 N205, KNG1 N169, ORM1 N56) indeed show the most pronounced increase in tri-antennary N-glycan species. Furthermore, we see that most other N-glycosites either had the same abundance of tri-antennary N-glycan species in both groups, or slightly higher abundance of tri-antennary N-glycan species in the pregnancy group. Only a single N-glycosite seemed to have a lower abundance of tri-antennary N-glycans (site 72 of ORM1), which also did not cluster with the tri-antennary dominant sites in pregnant women. Further inspection revealed that the tri-antennary N-glycans shifted towards tetra-antennary N-glycans, aligning with the trend of increased antennarity upon pregnancy. All in all, these sets of N-glycoproteins and N-glycosites define a well-defined and highly specific N-glycosylation signature for dissecting the pregnant group from the non-pregnant women.

Altogether, we may conclude that the N-glycoproteomes of mother and fetus are very different. Umbilical cord plasma displayed a relatively uniform N-glycosylation profile across most of the proteins and N-glycosites, characterized by the high abundance of fully sialylated di-antennary N-glycans. In contrast, pregnancy was associated with the increase of tri- and tetra-antennary N-glycans. These observations agree largely with previous N-glycomics (i.e. released-N-glycan) data comparing umbilical cord and maternal plasma^17,19^. Distinctively revealed by peptide centric N-glycoproteomics, we found that the increased branching was not uniform across all plasma N-glycoproteins, or even all liver-derived plasma proteins, but manifested itself primarily on a small subset of N-glycoproteins and even only on specific N-glycosylation sites. This directly illustrates the increased depth obtainable by peptide-centric N-glycoproteomics when compared to N-glycomics. The triad design was essential for distinguishing N-glycosylation features associated with pregnancy from those specific to fetal plasma. N-Glycosylation changed most drastically when comparing mothers to the umbilical cord plasma, which is not surprising as these individuals represent highly different developmental states. The non-pregnant women effectively provided a baseline to allocate certain features to either fetal or pregnant physiology.

### A subset of liver proteins display group-specific N-glycosylation features

We next sought to identify the proteins showing the largest group-specific shifts in N-glycosylation across the three study groups. Rather than comparing individual N-glycans separately, we again analyzed N-glycosylation at the N-glycosite level. This approach considers the complete heterogeneous N-glycan profile observed at a given site, thereby capturing changes in the overall N-glycosylation pattern rather than focusing on a single N-glycan structure. For each N-glycosite, we quantified the similarity between every pair of donors using cosine distance, similar to the earlier analysis. However, instead of comparing different N-glycosites within a donor group, we compared a single site between different donor groups. We then used an energy statistic to compare the similarity of profiles between donors within the same group to the similarity between groups. An illustrative example of this approach is shown in **Supplemental Figure 5**. This yielded a distinction score for each N-glycosite and each pairwise group comparison (pregnant women vs. control, pregnant women vs. fetus, and fetus vs. control), with higher scores (with a maximum score of 2) indicating greater group-specific differences in the overall N-glycosylation profile.

This revealed that, in addition to the proteins described earlier (KNG1 and ORM1), serum paraoxonase 1 (PON1), clusterin (CLU), and Immunoglobulin M (IgM) exhibited the most prominent altered N-glycosylation patterns on at least one N-glycosite, mostly distinctive between umbilical cord and the maternal plasma **(Supplemental Figure 6A)**. Due to the triad design, we were able to assign whether these altered N-glycoforms were fetal-specific (high similarity between non-pregnant control and pregnant women, e.g. CLU and IGHM), while others are pregnancy-specific (high similarity between non-pregnant controls and fetus, e.g. KNG1). Except for IgM (produced by B cells), the other proteins are predominantly expressed in the liver. In sharp contrast, a large set of abundant liver-derived proteins do not change N-glycosylation in either condition (e.g. C3, F2, HRG, TF), as is seen in **Supplemental Figure 6A**. This again showcases that the N-glycosylation changes are specifically observed in just a small subset of N-glycoproteins, most prominently PON1, KNG1, ORM1 and CLU.

Clearly site 253 shows no changes between groups, whereas changes in site 324 are very distinctive between all 3 studied groups. With the data used here, information on branching and exact sialic acid placement is unknown and therefore they are placed from left to right. **D)** Percentage of di- and tri-antennary N-glycans on site 324 of PON1. It was assumed that additional HexNAcs first branch and then extend. **E)** Summary of N-glycosylation changes observed on the four proteins presented here. The curved arrows annotate the rate-limiting enzyme for the conversion from one to the next N-glycan-category. The fetal signature is more towards di-antennary and hybrid-type N-glycans, while the pregnancy signature is more towards tri- and tetra-antennary N-glycans. Often, the non-pregnant women profile has both traits and represents seemingly an intermediate.

PON1 is a high-density lipoprotein (HDL)-associated antioxidizing enzyme synthesized primarily in the liver^28,29^. The protein abundance of PON1 is lower in the umbilical cord plasma, compared to both women groups, while pregnancy does not seem to affect the abundance of PON1 in plasma (**Figure 4A**). Both the N-glycosylation distinction score and the N-glycosylation profile of site N253 show identical N-glycosylation between groups, while site 324 shows a large distinctive difference in N-glycosylation landscape between all three studied groups (**Figure 4B**). The N-glycosylation landscape of the non-pregnant women has both di- and tri-antennary complex-type N-glycans, with varying degrees of sialylation. The most abundant N-glycan is N5H6S2, which can either be di-antennary with a longer antenna, or tri-antennary (**Figure 4C**). The umbilical cord plasma exhibits almost exclusively di-antennary complex-type N-glycans, in contrast to the pregnant women who exclusively display the larger N-glycans N5H6S2 and N5H6S3 (**Figure 4D**). It has been found that the activity of PON1 decreases as pregnancy progresses, both towards paraoxon and more drastically towards diazoxon^30,31^. As plasma concentrations seem to remain steady, we might hypothesize that the altered N-glycosylation found here affects PON1 activity, possibly due to steric interference or increased polarity around the N-glycosite, although further research should validate this hypothesis.

**Figure 4:**
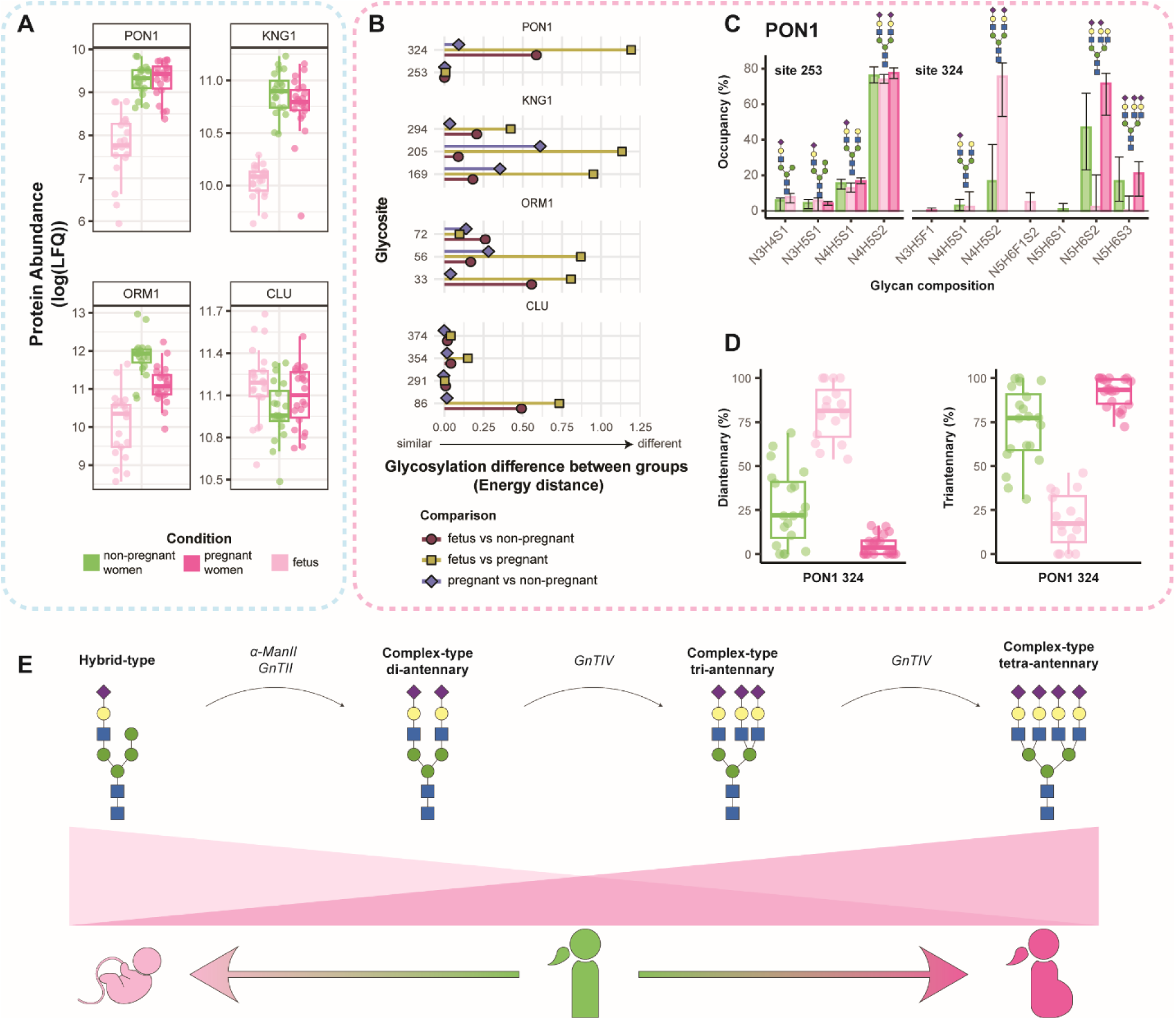
Liver proteins displaying the most distinctive N-glycosylation features for pregnant women and fetus. **A)** Protein abundance of serum paraoxonase 1 (PON1), kininogen 1 (KNG1), alpha-1-acid N-glycoprotein 1 (ORM1), and clusterin (CLU). The protein abundance is quantified as the log-transformed label-free quantification value, as determined by proteomics. Boxplots depict the median and the 25^th^ and 75^th^ percentiles. **B)** N-glycosylation change for each N-glycosite, calculated using the energy distance. Each dot represents a single comparison, where the red circle is fetus versus non-pregnant women, the yellow square is fetus versus pregnant women, and the blue diamond is pregnant versus non-pregnant women. A distance of 0 represents no difference and higher scores represent larger differences in N-glycosylation pattern between the groups. **C)** The N-glycosylation profile identified on PON1 on sites 253 and 324 in the different groups, where each bar represents the median occupancy and the error bars represent the spread from the 25^th^ to the 75^th^ percentiles. The major N-glycoforms are annotated with a N-glycan structure.

Another protein displaying altered N-glycosylation is KNG1, which is involved in the kallikrein-kinin pathway. KNG1 has been proposed as a putative biomarker for several hypertensive disorders in pregnancy, such as pregnancy-induced hypertension and pre-eclampsia^32^ . In our study, we found that the protein abundance is decreased in the umbilical cord plasma, while pregnant and non-pregnant women have about equal plasma concentrations of KNG1 (**Figure 4A**). However, the N-glycosylation on KNG1 is different between pregnant and non-pregnant women. All three N-glycosylation sites of KNG1 showed a shift towards the tri-antennary dominant cluster in **Figure 3**. Interestingly, the difference in antennarity between the non-pregnant and pregnant women is most pronounced on sites 169 and 205, while the difference between pregnant and non-pregnant is less pronounced on site 294, as also reflected by the distinction scores (**Figure 4B**, **Supplemental Figure 6B-C)**. On this latter site, the umbilical cord plasma shows a more distinctive N-glycoform landscape with a decreased abundance of larger N-glycans, while pregnant and non-pregnant women show more alike N-glycan signatures. Interestingly, all sites show variable degrees of fucosylation and sialylation, but pregnancy or infancy did not seem to affect this.

Alpha-1-acid N-glycoprotein (ORM1) is a negative immunomodulatory protein. In pregnancy, elevated levels of ORM1 have been associated with preeclampsia; together with increased levels in umbilical cord plasma ^33^. In our data, ORM1 abundance is lowest in umbilical cord plasma, and highest in non-pregnant women (**Figure 4A**). ORM1 has 5 N-glycosylation sites, of which we were able to cover three. The N-glycosite distinction score shows that site 33 is most distinct in the fetus, while site 56 is distinct in each group (**Figure 4B**). In non-pregnant women, all sites are covered with di- and tri-antennary complex-type N-glycans with varying degrees of sialylation and fucosylation **(Supplemental Figure 6D)**. Consistently, antennarity is decreased in umbilical cord plasma, and increased in the pregnant women **(Supplemental Figure 6E)**. Interestingly, again the degree of sialylation and fucosylation remained similar between the different groups.

The last protein we highlight showing a substantial N-glycosylation difference is clusterin (CLU). The overall protein concentration is similar between all groups (**Figure 4A**). CLU is a multi-functional protein: amongst others, it is involved in protein quality control, lipid metabolism through association with HDL, and complement regulation^34^. Elevated levels of CLU are also associated with preeclampsia^35,36^. CLU undergoes proteolytic cleavage, which results in the formation of two different chains^34^. Each chain has three N-glycosylation sites: sites 86, 103 and 145 are located on the beta chain, and sites 291, 354 and 374 are located on the alpha chain. These chains form a heterodimer, bound through disulfide bonds^34^. We cover sites 86 and 291 with multiple N-glycans in all three groups, while 354 and 374 were only consistently detected in umbilical cord plasma and pregnant women **(Supplemental Figure 6F)**. Site 86, located on the beta chain, specifically has altered N-glycosylation for the fetus, while both pregnant and non-pregnant women display similar N-glycosylation features (**Figure 4B**). Again, the difference was mainly the N-glycan antennarity: both women groups mostly have tri-antennary N5H6S3, while umbilical cord plasma has mainly N4H5S2. Sites 354 and 374 were also differently N-glycosylated between mother and fetus. Again, antennarity stood out as the main N-glycan trait; both sites had again higher occupancy of tri-antennary, fully sialylated N-glycans in the pregnant women **(Supplemental Figure 6G)**. Non-pregnant women mainly presented only a single N-glycoform, and other N-glycoforms appeared to be below the limit of detection, therefore stratifying whether these traits are pregnancy or infancy related is not possible. Interestingly, N-glycosite 291 did not change in any condition, again showcasing the distinct site-specificity of the altered N-glycosylation.

Altogether, these examples align well with a common trend that we observe: low antennary N-glycan species are relatively more abundant in umbilical cord plasma, while large and branched N-glycans are more abundant in pregnant women. The enzymes involved in the stratification between hybrid-type and complex-type N-glycans are α-mannosidase II (αMan-II) and N-acetylglucosaminyltransferase II (GnT-II), which consecutively remove mannose residues and attach the branching N-acetylglucosamine (GlcNAc) (**Figure 4E**). A third branch and fourth branch, which in plasma is most often a β1,4-linked branch, are added by N-acetylglucosaminyltransferase IV (GnT-IV). It seems that, in the fetal liver, these conversions happen less effectively, while in pregnancy the N-glycan machinery is more geared towards producing these tri-antennary and tetra-antennary N-glycans.

All previously discussed proteins are associated with HDL particles and/or the acute-phase response^37^, suggesting that the N-glycosylation remodeling may be related to changes in lipid metabolism and the slightly inflammatory state accompanying pregnancy. Interestingly, our proteomics analysis also revealed upregulation of proteins involved in protein-lipid complex remodeling and plasma lipoprotein particle remodeling (**Figure 2D**).

Cholesterol is essential for fetal development. Whereas low-density lipoproteins are the main carriers of cholesterol in adults, HDL has a more prominent role in fetal cholesterol transport^38^. In addition to endogenous synthesis, the fetus can acquire maternal cholesterol through the placenta. Lipid metabolism is also substantially altered during pregnancy. During the last trimester, maternal metabolism shifts toward a more catabolic phase, characterized by reduced fat accumulation and decreased insulin sensitivity, while HDL particles increase in size and undergo compositional changes compared to non-pregnant women^39^. Fetal and maternal HDL also differ in composition; ApoE is enriched in fetal HDL, while all other apolipoproteins are decreased^40^.

PON1 provides an example of the relationship between lipid metabolism and N-glycosylation. PON1 is a calcium-dependent multifunctional enzyme: amongst others, it has paraoxonase, arylesterase and lactonase activities; it is a component of HDL particles in circulation; it reduces low-density lipoprotein (LDL) oxidation; and it reduces macrophage foam cell formation^28^. The activity of PON1 is influenced by ApoE and ApoA-1, which are the major proteins in HDL in fetus and adult, respectively^40,41^. In the fetus, both the abundance and the activity of PON1 are decreased^42^. In pregnancy, the enzymatic activity is also decreased^30,31,43^, despite unchanged plasma abundance. We observe a strong shift to exclusively tri-antennary N-glycan species in pregnancy. Fully sialylated tri-antennary N-glycans are larger and more polar compared to their di-antennary counterparts, which might negatively influence the HDL incorporation or function. In addition to PON1, both CLU and KNG1, as well as the previously discussed SERPINA1, SERPING1 and VTN, can associate with HDL^37,39^ and showed similar substantial changes in N-glycosylation. These findings raise the possibility that pregnancy-associated changes in HDL metabolism are accompanied not only by changes in particle composition and protein activity, but also by selective remodeling of their protein N-glycosylation.

### Fetal IgM is distinctively N-glycosylated hinting at substantial fetal production

Another protein displaying a substantial N-glycosylation shift in our data was IgM, which belongs to the larger family of immunoglobulins. During pregnancy and in the early weeks of infancy, only IgG antibodies, the most abundant isotype in adult blood, are actively transported over the placenta by the neonatal Fc receptor (FcRn)^44,45^. This provides the fetus and newborn with the adaptive humoral immunity repertoire of the mother. It is thus generally assumed that the fetus/infant barely produces its own antibodies^46^. Therefore, we examined the immunoglobulins in detail, starting with their abundances (**Figure 5A**). When comparing the IgG subclasses, we noticed that each of the subclasses had a slightly higher abundance in the umbilical cord plasma, compared to both adult groups. Interestingly, the ratio of the subclasses was also slightly different: umbilical cord plasma seems to contain relatively more IgG1 than IgG2, whereas the relative abundances of IgG3 and IgG4 were similar across all sample groups (**Figure 5B**). This is in line with previous reports stating a higher transport rate of IgG1 and a lower transport rate of IgG2 over the placenta^47–49^. N-Glycoproteomics further revealed that IgG1 and IgG2 have very similar N-glycan characteristics across the mother and umbilical cord pairs (**Figure 5C**).

**Figure 5:**
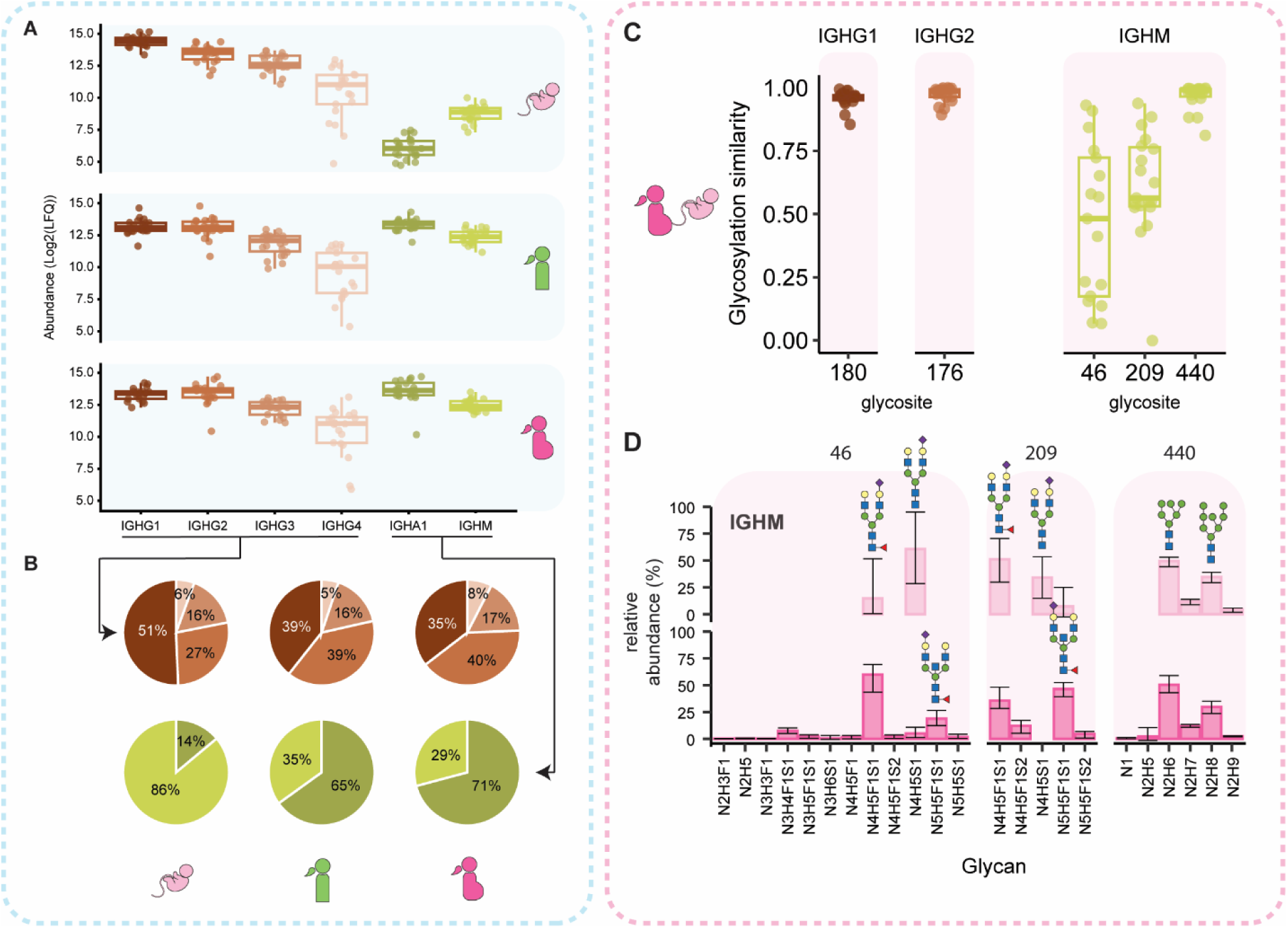
Fetal IgM is distinctively N-glycosylated. **A)** Abundance (quantified as the log-transformed label-free quantification value) of different immunoglobulins in fetus (top panel), non-pregnant women (middle panel) and pregnant women (bottom panel). Most notably, IgA1 is about 1C0-200 times lower, while IgM is only about 10 times lower in umbilical cord plasma than in maternal plasma. Boxplots depict the median and the 25^th^ and 75^th^ percentiles. **B)** Pie charts depicting the composition of IgG subclasses (top) and IgA and IgM (bottom), for fetus (left), non-pregnant women (middle) and pregnant women (right). In the fetus there is relatively more IgG1, and less IgG2. Furthermore, there is relatively high abundance of IgM in umbilical cord plasma compared to IgA1. **C)** N-glycosylation similarity of matched mother and fetus samples for each N-glycosite, where 1 is highly similar and 0 is dissimilar. Boxplots depict the median and the 25^th^ and 75^th^ percentiles. Individual datapoints are represented as dots. **D)** Bar plot of median N-glycan compositions on IgM. The bars represent the median relative N-glycan abundance, and the error bars represent the spread from the 25^th^ to the 75^th^ percentiles. The major N-glycoforms are annotated with a N-glycan structure.

Next to IgG, IgA1 and IgM were quantified across all samples. IgA2 and IgD were also quantified in the adult donors, but were too low abundant to be detected in the fetus, in agreement with later development of fetal humoral memory^46,50^. IgA1 levels were detectable, albeit about 160-200-fold less abundant in umbilical cord plasma compared to the adult groups, as expected, since IgA cannot be transported over the placenta. However, although IgM was also lower in umbilical cord plasma compared to maternal plasma, the fold change was just 11-fold (**Figure 5B**). As there are no known placental transport mechanisms of antibodies besides FcRn-mediated transport of IgG, which does not bind other isotypes, the relative moderate abundance of IgM in the fetus suggests production of endogenous fetal IgM already at birth. It has been reported that fetal B-cells can produce IgM before birth^51,52^. And although the IgM repertoires have been found to be already diverse at birth, the low frequency of somatic mutations suggests that IgM production is antigen-independent^53,54^. This already shows that fetal IgM is different from adult IgM.

Strikingly, we observed that the N-glycosylation characteristics on IgM were very clearly distinct between fetus and the mothers (**Figure 5C-D**). We were able to cover 3 of the 5 N-glycosylation sites on IgM. The N-glycosylation on site N440 in the C-terminal tail-piece did not substantially change between the two groups. However, the sites N46 and N209 showed very distinct N-glycosylation patterns in the fetus compared to the mothers. The N-glycosylation profile of fetal IgM had less fucosylation on site N46 and less bisection and less fucosylation on site N209, compared to IgM observed in pregnant women. We also compared the N-glycosylation of IgM to that of the control women, which was more alike the pregnant mothers, indicating that we observe specific features indicative of fetal IgM (**Supplemental Figure 7**).

We propose that the markedly distinct N-glycosylation of IgM in umbilical cord plasma, primarily at two N-glycosylation sites, provides an additional molecular feature distinguishing fetal from adult IgM. The IgM repertoire at birth is different from adult IgM: the limited frequency of somatic mutations in fetal IgM represents a more germline-encoded repertoire, whereas adult IgM is shaped by the adaptive immune system^53,54^. Together with the distinct N-glycosylation we found on fetal IgM, these differences suggest that fetal IgM is produced and processed differently from adult IgM. In line with this, previous studies showed that the antigen-binding specificity of fetal IgM differs between matched maternal and umbilical cord pairs^55^. A possible explanation is that fetal IgM represents ‘natural antibodies’, produced by B-1 cells, in contrast to the conventional B-2 cells, which are most abundant in adults^56^. There are many differences between B-1 and B-2 cells, including their localization, surface receptors and response to stimuli^56,57^. These differences may result in distinct N-glycosyltransferase activities and consequently different N-glycan profiles, as observed here. Whether these N-glycosylation differences reflect the distinct developmental origin of fetal B cells, differences in N-glycosyltransferase activity, or the antigen specificities encountered in utero remains to be established.

Finally, the moderate but notable level of IgM production in utero is distinctive from IgA (**Figure 5A and 5B**), which seems only marginally present in umbilical cord blood. As both these immunoglobulin moieties are likely equally hard to be transferred across the placental barrier it underscores the unique favored production of fetal IgM.

### Conclusions and limitations

Here, we performed proteomics and site-specific N-glycoproteomics on a cohort of matched maternal and umbilical cord plasma, which represent pregnant and fetal physiology, respectively. We supplemented this cohort with a control group of non-pregnant women, to enable us to assign molecular features specifically to pregnancy or fetal physiology. To our knowledge, this is the first study to quantify plasma N-glycosylation in a site-specific manner across pregnant women, paired umbilical cord plasma, and healthy non-pregnant women. The data demonstrate that both fetal plasma and pregnancy are characterized by distinct proteomics and N-glycoproteomics signatures. Both pregnancy and fetal physiology are characterized by changes in proteins involved in complement activation. Furthermore, pregnancy was associated with selective, site-specific remodeling of N-glycosylation on a defined subset of liver-derived N-glycoproteins, suggesting targeted regulation rather than a global N-glycan shift. Most of the proteins with altered N-glycosylation were involved in protein-lipid particles. Lastly, in contrast to IgA, IgM was found moderately abundant in the umbilical cord plasma and carried distinct site-specific N-glycosylation profiles on two N-glycosites, adding a N-glycoproteomics dimension to the distinction of fetal IgM from adult IgM. The protein- and site-specific increase in N-glycan branching during pregnancy, together with the fetal-specific N-glycosylation of IgM, illustrates the additional biological information obtained by resolving N-glycosylation in a protein- and site-specific manner.

Although we carefully designed our study, several limitations can still be considered. First, all serum samples were obtained at a single time point, preventing assessment of how individual proteins and N-glycosylation sites change throughout gestation. Longitudinal sampling across pregnancy would provide deeper insight into the timing and regulation of these N-glycosylation changes and could help distinguish early adaptations from changes associated with late gestation. Second, the study focused on healthy, uncomplicated pregnancies and therefore cannot determine whether the observed N-glycosylation patterns are altered in pregnancy complications. Future studies incorporating longitudinal sampling and disease cohorts, particularly preeclampsia and other disorders of maternal-fetal physiology, may establish whether site-specific plasma N-glycosylation can provide mechanistic insight into disease or serve as a marker of altered pregnancy physiology. Lastly, in the analysis we focused on the most abundant proteins, to minimize missing values and ensure robustness and quantification with statistical significance. Deeper (N-glyco)proteomic profiling, using for example fractionation or additional enrichment strategies, may reveal additional protein and N-glycosylation signatures.

## Methods

### Study cohort

The matched umbilical cord-maternal plasma samples used in this study were collected directly after delivery (average gestation time 37.9 weeks). All women had an uncomplicated pregnancy and neonatal outcomes for all children were optimal. Signed informed consent was obtained from all women, and the collection of blood samples and clinical data received approval by the Ethics Committee of the Leiden University Medical Centre (P02-200). These plasma samples have been described and studied previously^16^. There were 22 maternal plasma samples and 21 umbilical cord plasma samples, for two maternal plasma samples we did not have a matched umbilical cord plasma sample and one mother gave birth to twins. Details on the matched maternal and umbilical cord plasma cohort can be found in **Supplemental Table 1.** Plasma samples of non-pregnant women were obtained from the Sanquin National Screening laboratory. The donors’ ages ranged from 20 to 36 years (median age ∼31). These samples have also been studied previously^58^. More details of the non-pregnant control samples can be found in **Supplemental Table 2.**

### Sample preparation

5 µL of plasma was diluted in 80 volumes reduction and alkylation buffer (Tris HCl (100 mM), TCEP (40 mM), CAA (160 mM); pH 8.5), and proteins were denatured at 80 °C for 10 minutes. Proteins were digested with trypsin (1:100) and LysC (1:100) overnight at 37 °C. Peptides were desalted using Oasis PRiME HLB (10 mg) 96-well plates (Waters). N-Glycopeptides were enriched using cotton-HILIC SPE, as described previously^59^. The flow-through was collected for proteome analysis and the enriched fraction was used for N-glycoproteomics. The entire sample preparation was performed in 96-well plates, samples were plated randomly, and pipetting was automated on the Bravo liquid handling system (Agilent Technologies).

### LC-MS/MS analysis

Peptides were resuspended in 0.1% TFA and analyzed using an Ultimate 3000 HPLC system (Thermo Fisher Scientific) coupled online to a ZenoTOF 7600 system mass spectrometer (SCIEX), equipped with the OptiFlow Nano Source. Peptides were trapped on a PepMap Neo Trap Cartridge heated at 32 °C (5 μm, C18, 300 μm x 5 mm, Thermo Fisher Scientific) and separated using a 15 cm IonOptics Aurora Elite XS (75 μm ID, 1.7 μm, C18) analytical column. Buffer A was 0.1% formic acid in water, buffer B was 0.1% formic acid in 80% ACN. For the proteomics, a 65-minute method was used: starting at 4% buffer B to 9% B in 2 min, from 9% to 30% in 37 min, from 30% to 44% in 15 min, from 44% to 55% in 4 min, from 55% to 99% in 1 min, 99% wash-out in 5 min and re-equilibration back to 3% buffer B in 10 min.

For the N-glycopeptides, a 135 min method was used: starting at constant 3% buffer for 1 minute, ramping to 10% B in 1 min, from 10% to 35% in 30 min, from 35% to 55% in 4.5 min, from 55% to 99% in 0.5 min, 99% wash-out in 4.5 min and re-equilibration back to 4% buffer B in 10 min.

The proteomics data was recorded in data-independent fashion, using ZT SWATH with default settings while estimated LC PWHH was set to <=4.5 s. N-Glycoproteomics data was recorded in information dependent acquisition, with the parameters optimized previously^59^. Briefly, the MS^1^ range was set to 700-2000 m/z, with an accumulation time of 50 ms and a CE of 8 V. The top 15 precursors (charge 2-12) were selected for fragmentation with EAciD (EAD: 9 eV, CID: dynamic CID enabled with slope of 0.049 and intercept -11 and slope of 0.050 and intercept -17 for charges 2+ and 3+ and higher, respectively), and dynamic exclusion was set to 12 s. The acquisition and reaction time were set to 65 and 20 ms, respectively.

### Data analysis

Bulk proteomics data was processed using DIA-NN v.1.9 in library-free mode^60^. 1% false discovery rate (FDR) was used for precursor filtering, and trypsin/P was used as protease with maximum two allowed missed cleavages. The default cysteine carbamidomethylation and methionine oxidation were used as modifications, with the maximum number of variable modifications set to 1. Mass accuracy was determined separately for each run, and the quantification method was set to legacy. The heuristic protein inference was disabled while the unrelated runs, isotopologues, and MBR were enabled. A blood protein database previously described was used for the data search^20^. Briefly, this database consists of 2,445 common blood proteins according to the Human Protein Atlas^61^ (proteinatlas.org, accessed in November 2023), and their 5% most frequent single amino-acid variations acquired from NextProt database (accessed in May 2024^62^). Immunoglobulin sequences and common umbilical cord plasma proteins were manually added to the database (see Resource availability). The complete report generated from DIA-NN was used for further downstream analysis. The protein sequences describing the variable domains of immunoglobulins and proteins identified with fewer than 2 peptides were removed from the dataset. Contaminants like keratins, platelet proteins and erythrocytes were also removed from the dataset. Proteins detected in at least 10 samples and within at least one of the three groups (non-pregnant women, pregnant women, umbilical cord plasma) were kept for further downstream analysis. The protein abundance (MaxLFQ) was calculated using the diann_maxlfq function of the DIAgui package (https://github.com/mgerault/DIAgui), with Q.Value, Lib.Q.Value and Lib.PG.Q.Value, all set at 1% threshold. Missing values were imputed using random values between 50% and 100% of each protein’s minimum observed abundance. Throughout this manuscript, proteins are referred to by their gene name.

N-glycoproteomics data was analyzed using MSFragger (Fragpipe version 23.1; MSFragger version 4.4.1; IonQuant version 1.11.20) using a modified N-glyco-LFQ workflow with semi-specific cleavage for trypsin/P and match-between-run enabled^63,64^. The same protein database as for the proteomics data was used. Workflow file and N-glycan database are deposited in the MassIVE repository. A 1% FDR threshold was used for filtering of N-glycoproteomics identifications, and MaxLFQ values were used for quantification. Further PSM filtering was based on protein haplotypes detected in each individual: i) haplotype (protein isoform variations) was defined for each protein in each donor if supported by at least three different (modified) peptides ii) data was grouped by haplotype and N-glycosite; iii) peptide sequences (e.g. semi-specific sequences, missed cleavages) were filtered to be found in at least 80% of donors with the same haplotype; iv) unique modified peptides (sequence + N-glycan + other modifications) were filtered to be found in at least 50% of the donors with the same haplotype and donor group (i.e. pregnant, non-pregnant, or cord). In addition, peptides containing more than one potential N- or O-N-glycosylation site according to UniProt were excluded because the available fragmentation evidence and chromatographic separation was not sufficient for reliable modification localization. Frequently misassigned N-glycan compositions were further evaluated by manual inspection of the corresponding peptide-spectrum matches (PSMs) and reassigned where necessary. N-Glycopeptide abundances were summed per N-glycosylation site, and used to compute relative abundances. Antennarity was inferred from the N-glycan composition, where it was assumed that additional GlcNAcs first branch and then extend, as shown in **Supplemental Figure 5 C, E, G)**.

For each donor and N-glycosylation site, the N-glycan profile was represented as a vector of relative N-glycan abundances. The dissimilarity between two site-specific N-glycan profiles, (X) and (Y), was quantified using cosine distance:

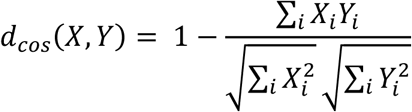

Recently proposed compositional-data transformations^65^ were not used because they showed low robustness to poorly quantified low-abundance N-glycoforms.

To identify N-glycosylation sites exhibiting systematic differences between cohorts, an energy distance was calculated between the cohort-specific distributions of N-glycan profiles, using cosine distance as the underlying pairwise dissimilarity. For cohorts containing profiles (X_1, …, X_n) and (Y_1, …, Y_m), respectively, the distance was defined as

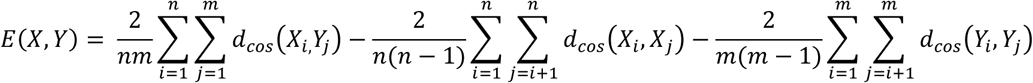

For within-cohort clustering, cosine distances between every pair of N-glycosylation sites were calculated separately for each donor. The median donor-level distance was then used as the pairwise distance between the two sites. A low value therefore indicated that the sites displayed similar N-glycan compositions consistently across donors.

To focus the clustering analysis on heterogeneous sites for which changes in N-glycan composition could be meaningfully resolved, sites whose median fraction of the most abundant N-glycan exceeded 0.90 were excluded. Average-linkage agglomerative hierarchical clustering was then performed using the resulting pairwise distance matrix, and clusters were defined by cutting the dendrogram at a distance threshold of 0.35.

Statistical tests were performed using the Kruskal-Wallis rank-sum test and Kruskal-Wallis Multiple Comparisons using the stats (v 4.4.0) and FSA (v 0.10.0) packages. Multiple testing correction was performed using the Benjamini-Hochberg method. Plots were made using the following packages: ggplot2 (v 3.5.1), ggpubr (v 0.6.0), ggrepel (v 0.9.5), heatmaps were made using ComplexHeatmap (v 2.20.0). Other used packages are tidyverse (v 2.0.0) and protti (v 0.9.0).

## Resource availability

### Lead contact

Albert J.R. Heck

### Data and Code availability

The raw LC-MS/MS data files, protein and N-glycan databases, the DIA-NN and MSFragger output, and the processed data files generated in this study have been deposited in the MassIVE repository under the identifier MSV000103405 and password Reviewer_y4deCK

### Author Contributions

**S.J.**: conceptualization, methodology, formal analysis, investigation, writing - original draft.

**S.K.N.:** methodology, formal analysis, investigation, writing - original draft. **G.R.**: data

analysis, writing – reviewing & editing. **F.R.I.**: methodology, writing – reviewing & editing. **R.V.**: resources. **P.S.:** supervision, writing – reviewing & editing. **G.V.:** resources, writing – reviewing & editing. **A.J.R.H.**: conceptualization, supervision, writing - original draft, funding acquisition.

## Acknowledgements

This project was made possible through the Large-Scale Research Infrastructure - National Roadmap consortia BioBeyond_NL (grant number 184.037.015, Grant ID: https://doi.org/10.61686/IONOX39631) and PICTURE (grant number 184.2025.001) of the Netherlands Organisation for Scientific Research (NWO). A.J.R.H. acknowledges support from NWO through the Spinoza Award SPI.2017.028.

