## Supplemental Figures 1-7, Supplemental Tables 1-2 for "Disentangling maternal pregnancy and fetal-specific plasma signatures through integrated proteomics and site-specific N-glycoproteomics"


| *Title* | *Page* |
|---|---|
| *Supplemental Figure 1* | *2* |
| *Supplemental Figure 2* | *3* |
| *Supplemental Figure 3* | *5* |
| *Supplemental Figure 4* | *6* |
| *Supplemental Figure 5* | *7* |
| *Supplemental Figure 6* | *8* |
| *Supplemental Figure 7* | *10* |
| *Supplemental Table 1* | *11* |
| *Supplemental Table 2* | *12* |
| *Supplemental Table 3* | *xlsx* |

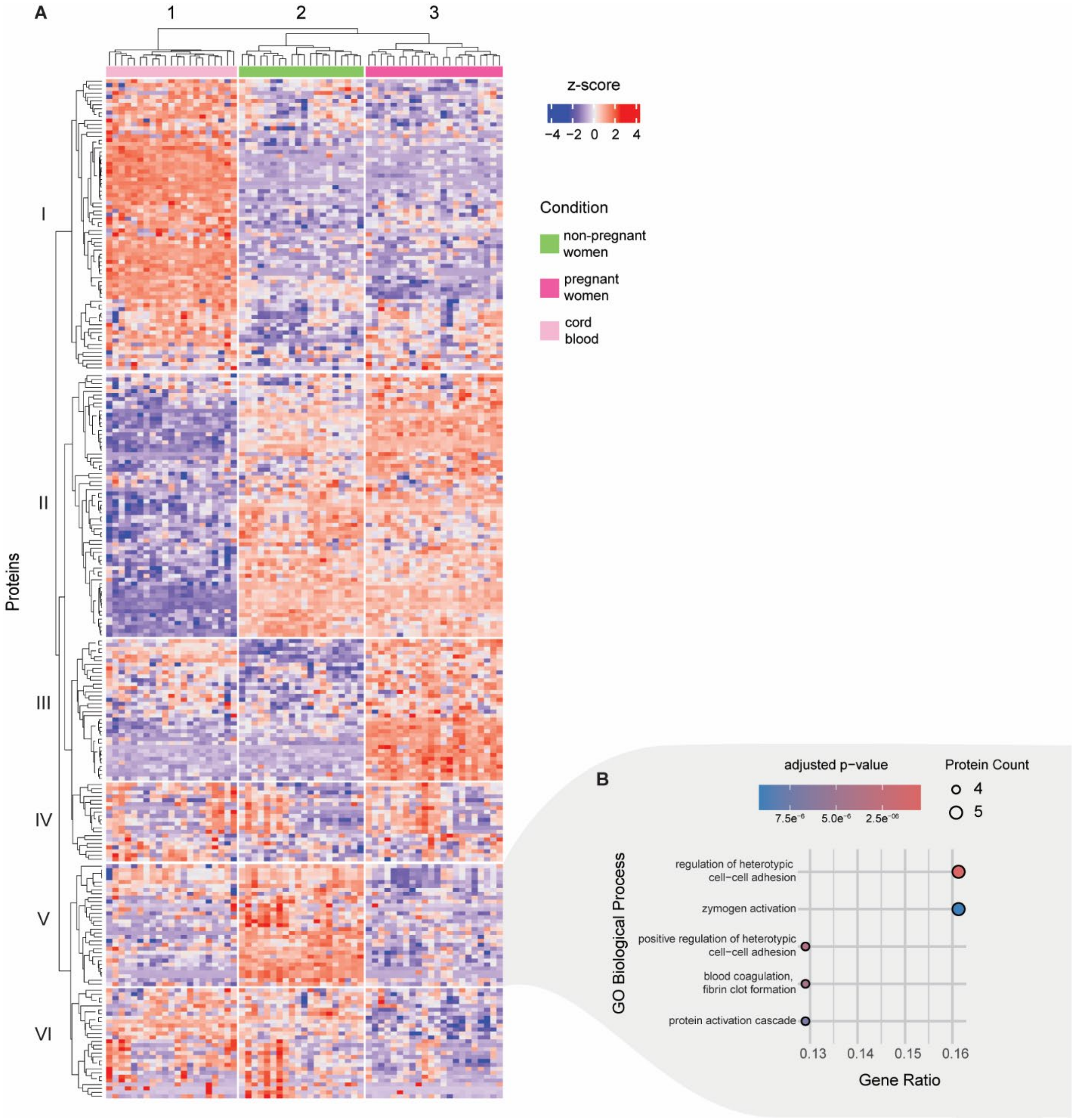


***Supplemental Figure 1: Proteomics: complete quantitative heatmap of all detected proteins. A)*** *Hierarchical clustering shows clear clusters of the three groups (non-pregnant women, pregnant women and fetus (umbilical cord plasma)). Cluster I to III are described in the main Figure 2. Cluster IV, V and VI contain 20, 31, and 28 proteins, respectively. An additional cluster, cluster V, has several proteins downregulated in both pregnant women and fetus.* ***B)*** *GO enrichment analysis of cluster V proteins. Cellular adhesion, zymogen activation and coagulation are amongst these terms, which suggesting alternative regulation in both mother and neonate. All underlying proteomics data is available via Supplemental Table 1*

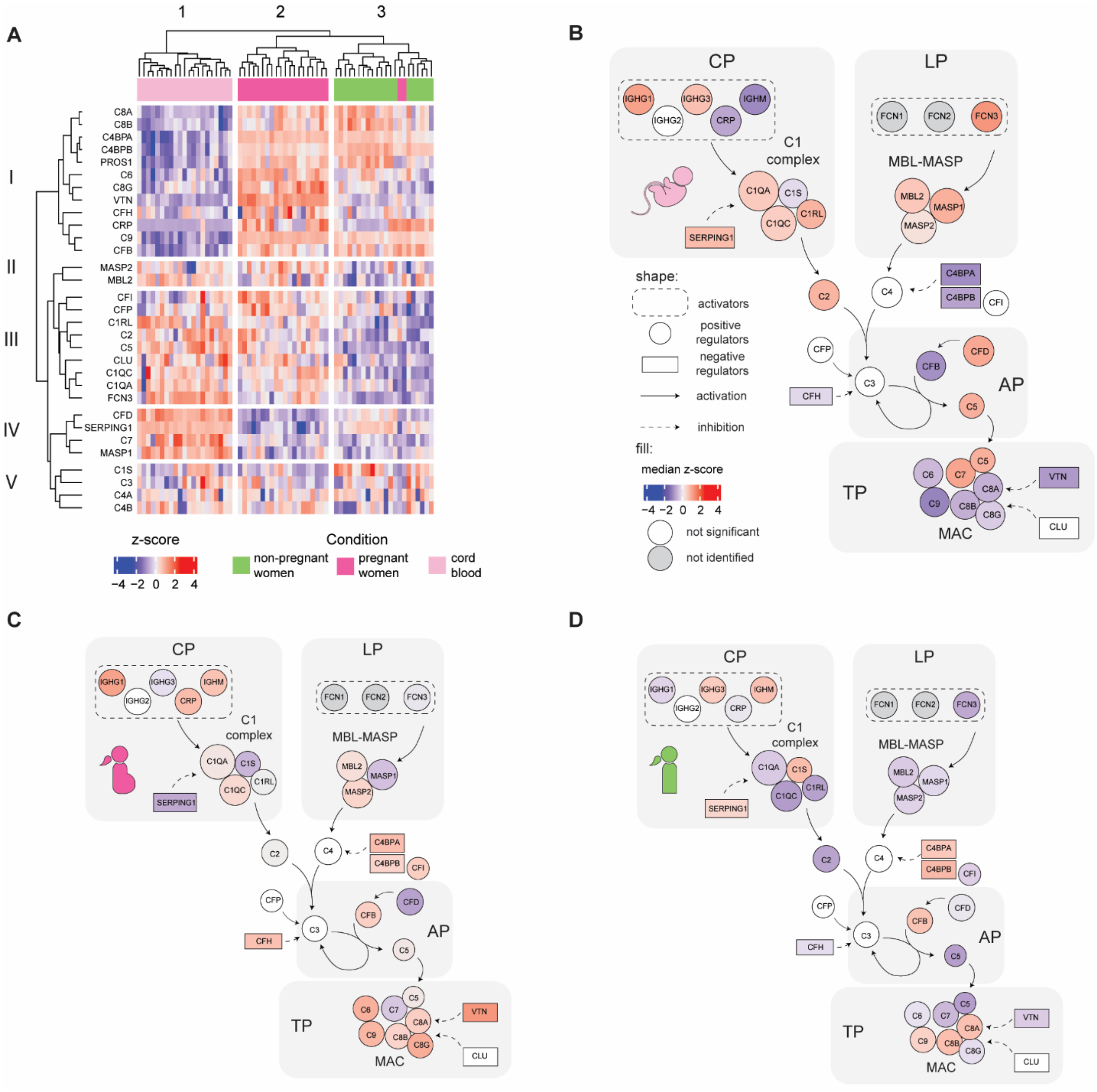


***Supplemental Figure 2: Complement rearrangement in fetus and during pregnancy. A)*** *Selected heatmap of exclusively proteins involved in the complement system. Again, hierarchical clustering separates fetus from both pregnant and non-pregnant women. Also, pregnant and non-pregnant women separate, albeit less prominent. In umbilical cord plasma, proteins in cluster I are downregulated. This cluster contains most of the complement inhibitors, as well as the complement proteins involved in the MAC-complex. Both fetus and pregnant women have high abundance of proteins in cluster III, containing proteins initiating the classical pathway (CP) and lectin pathway (LP); suggesting increased complement activation in both groups. The complement cascade is shown schematically in* ***B, C, and D****, where each circle represents a protein and the color represents the median z-*

*score normalized abundance of that protein in fetus (B), pregnant women (C), and non-pregnant women (D). Grey colored proteins were not identifiable and/or quantifiable, and white colored proteins were not significantly different between the groups (after Kruskal-Wallis rank sum test).*

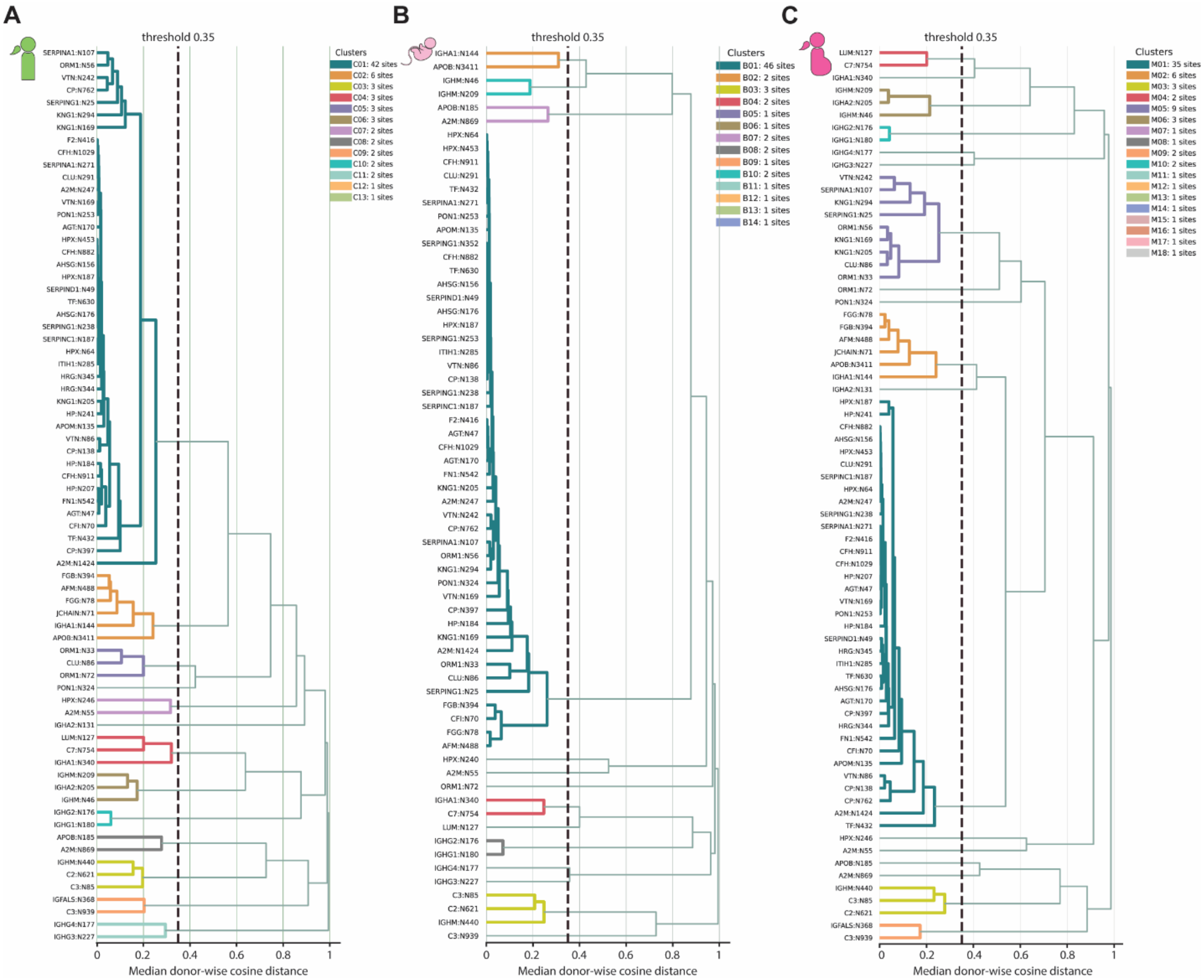


***Supplemental Figure 3: N-glycosite clustering based on cosine distance.*** *Dendrograms of N-glycosite clustering in **A)** non-pregnant women, **B)** fetus, and **C)** pregnant women. Glycosites were clustered based on the cosine distance of the quantitative glycan distribution. A threshold of 0.35 was used to define separate clusters, which are color coded. Gene name and glycosite number are indicated along the y-axis. Glycosites that cluster together present with similar glycan profiles. In all groups, the glycosites in the largest cluster (cluster 1 in blue) are mainly dominated by bi-antennary complex-type glycans. Furthermore, glycosites of IgM (440), C2 (621) and C3 (85) cluster together, which are known to be glycosylated with mainly oligomannose type glycans (cluster 3 in yellow).*

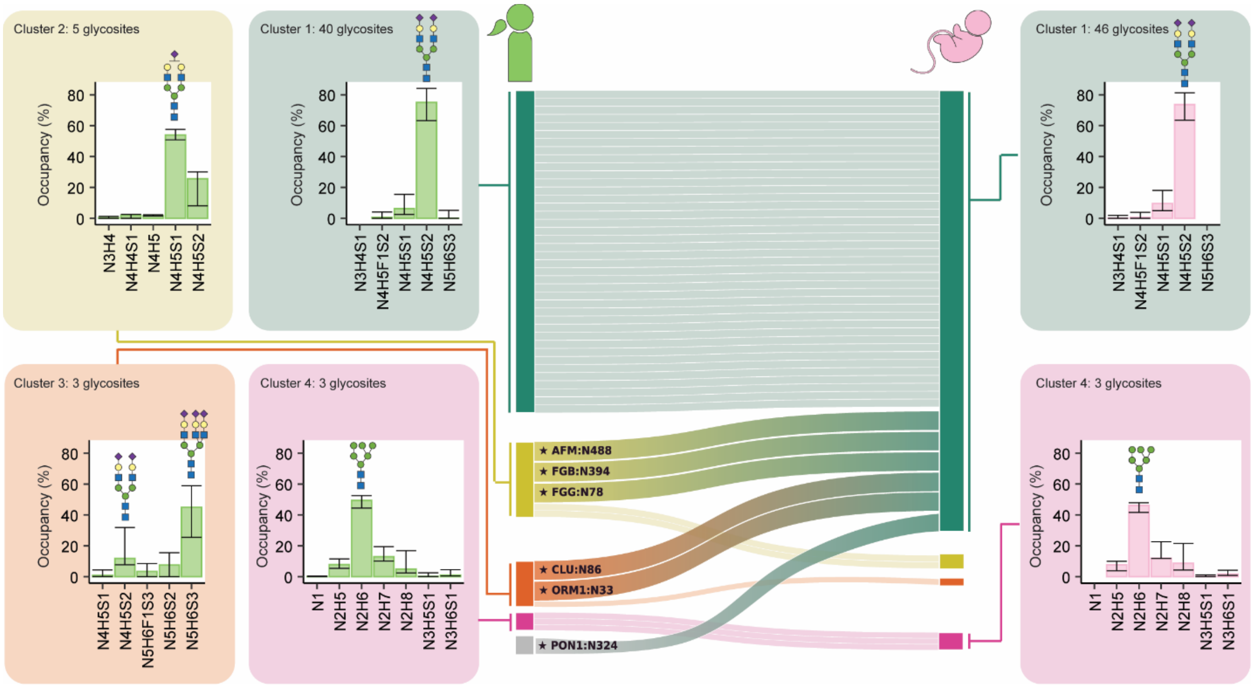


***Supplemental Figure 4: N-glycosite profiles shift distinctively especially towards N4H5S2-dominant in umbilical cord plasma.*** *Sankey diagram displaying how N-glycosites cluster based on glycosylation similarity, only clusters with 3 or more glycosites are shown here. Each line represents a single N-glycosite. Four major clusters with multiple N-glycosites are classified in non-pregnant women, while only two major clusters exist in the fetal plasma. Several glycosites that belonged to cluster 2 and 3 (glycosylation dominated by N4H5S1 and N5H6S3, respectively), cluster with glycosites in cluster 1 in the fetus (glycosylation dominated by N4H5S2). The N-glycosites that move between clusters are annotated with gene name and glycosite number. On the sides, the average glycan distribution across all glycosites in the different clusters is shown. Error bars represent the 25th and 75th percentile, and the major glycoforms in the cluster are annotated using cartoons.*

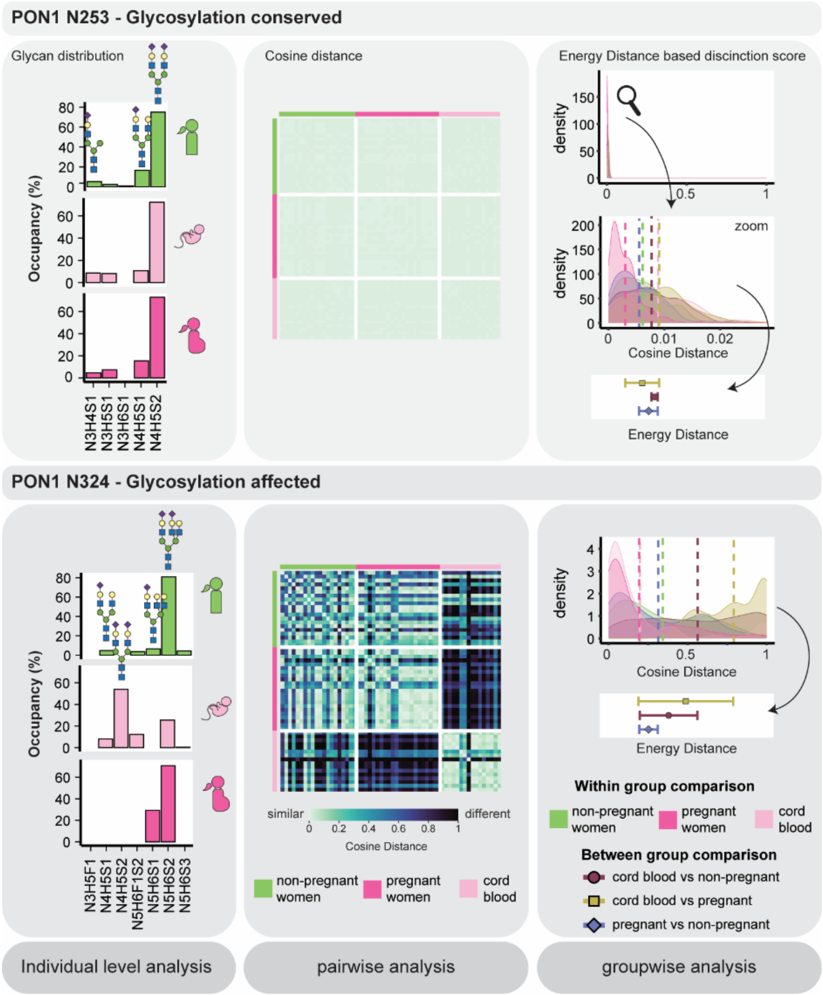


***Supplemental Figure 5: Multi-type analysis comparing N-glycosites.*** *Depicted is the approach used to compare glycosylation profiles between individuals and groups. In the example, we look at a conserved glycosylation site (similar between groups and individuals) on top and at a glycosylation site affected by the different physiologies of the donors (different glycosylation between groups) at the bottom. The analysis has 3 levels, indicated from left to right: the individual glycan level, a pairwise comparison, and a groupwise comparison. First the glycosylation profile is examined for each individual (individual glycan level, left panel). The analysis focusses on which glycans are present and in which quantities. Next, these glycan profiles are compared between donors in a pairwise analysis using cosine distances; this reveals which individuals have similar glycosylation (middle panel). The cosine distance is scaled from 0 to 1, where 0 represents similar profiles and 1 represents completely different profiles. Then, the within group similarity is compared to the between group similarity (groupwise analysis, right panel). The differences between these two factors determines is the 'energy distance', which reflects how different a glycosite is between two groups. The maximum energy distance is dependent on the standard deviation*

*within groups and the distance between two groups, it ranges from slightly negative values to a maximum of 2. The exact formulas can be found in the Methods section.*

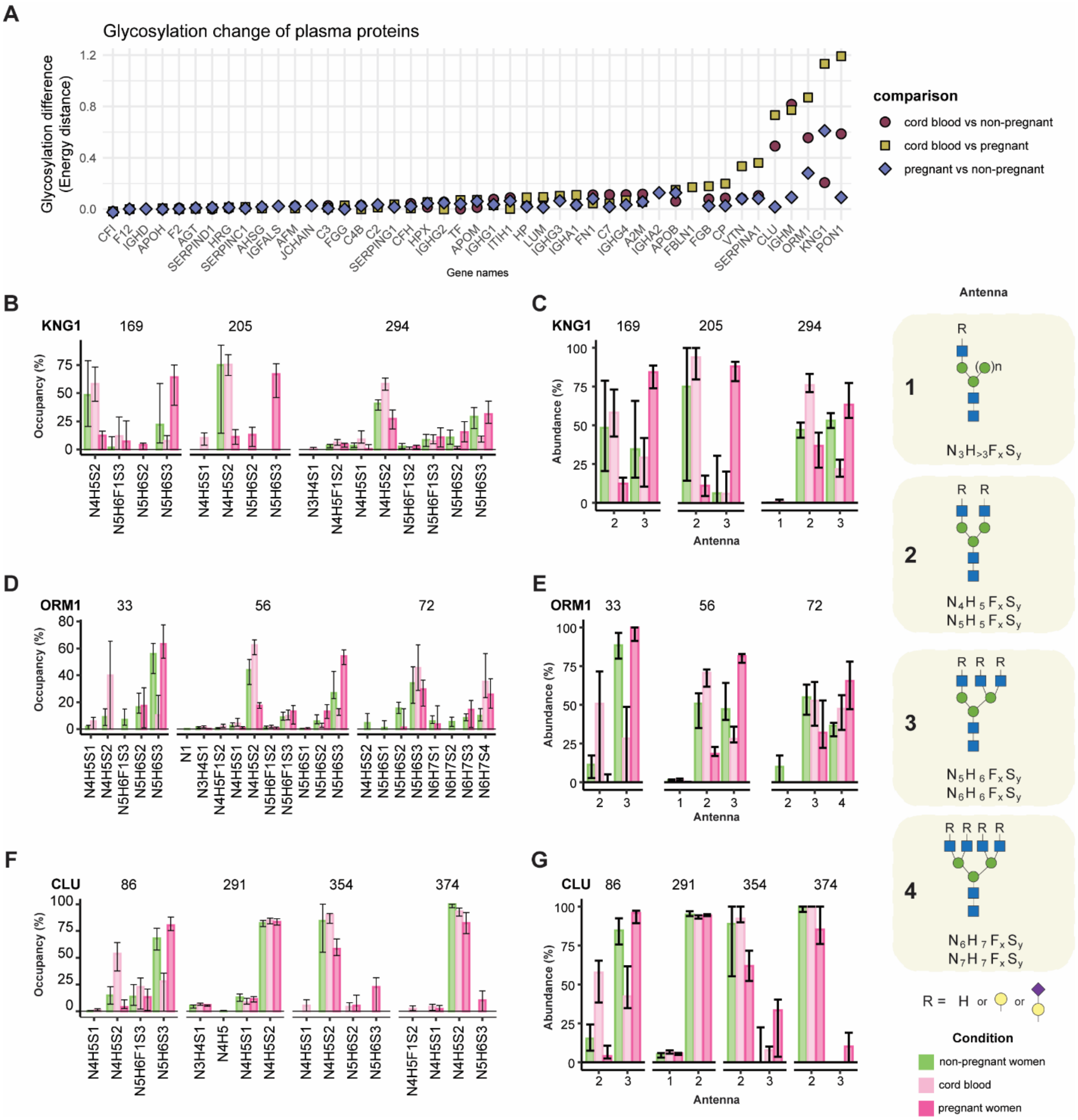


***Supplemental Figure 6: Most prominent N-glycosylation changes in plasma proteins. A)*** *The maximum glycosylation change for each glycoprotein across glycosites, calculated using the energy distance. Each dot represents a single comparison, where red circle is fetus versus non-pregnant women, yellow square is fetus versus pregnant women, and blue diamond is pregnant versus non-pregnant women. A large energy distance represents a large difference in glycosylation pattern between groups. Most proteins do not change drastically in their N-glycosylation, while a selected subset does undergo a larger change on at least*

*one glycosite.* ***B, D, and F)*** *The N-glycosylation profile identified on KNG1 (B), ORM1 (D), and CLU (F) in the different groups, where each bar represents the median occupancy and the error bars represent the spread from the 25$^{th}$ percentile to the 75$^{th}$ percentile. The number above each panel represents the glycosite.* ***C, E, and G)*** *Bar graph translating the glycosylation profile to an antenna distribution for KNG1 (C), ORM1 (E), and CLU (G). The number above each panel represents the glycosite. The classification of the antenna is described on the left with a glycan cartoon and a composition formula where x and y are any number from 0 to the number of antenna. In B-G, the color of the bar represents each group, where green is non-pregnant women, light pink is fetus and dark pink is pregnant women. We observe a consistent shift towards increased antennarity in pregnant women.*

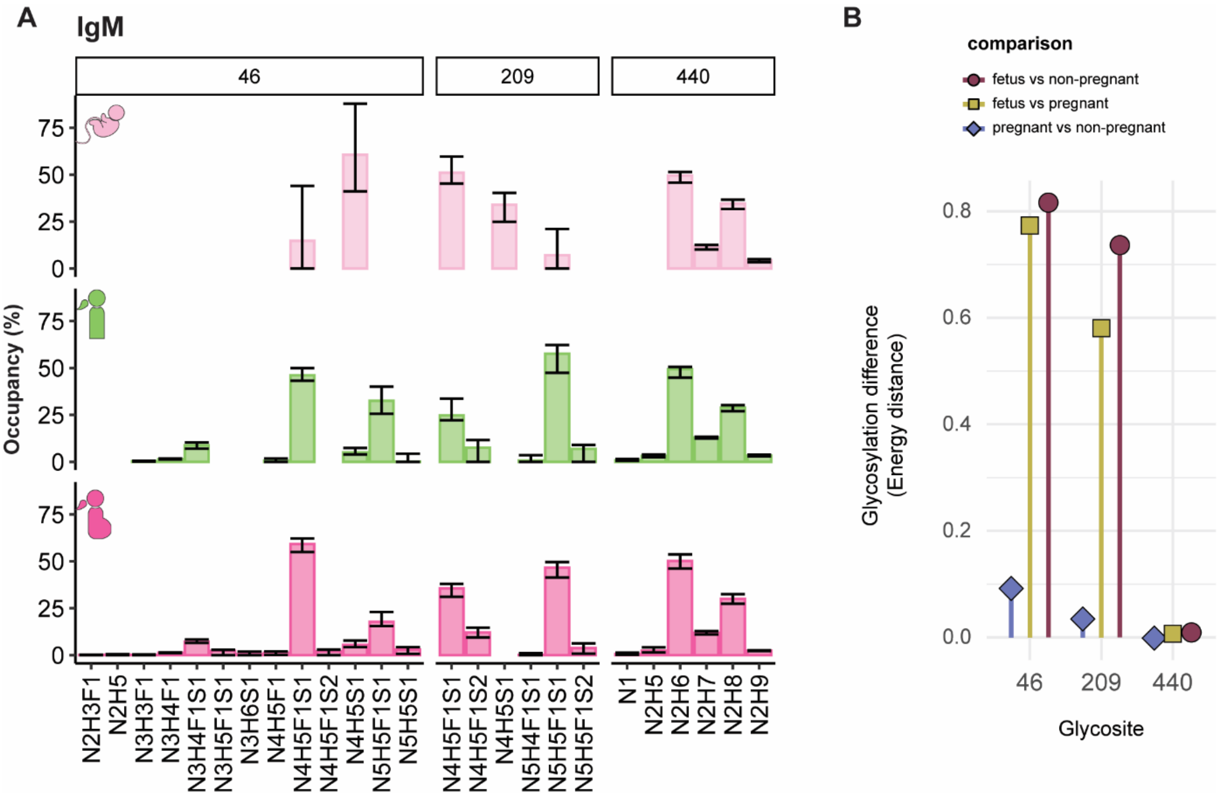


***Supplemental Figure 7: Umbilical cord plasma IgM exhibits an independent unique glycosylation signature. A)*** *Bar plot of median glycan compositions observed on IgM in umbilical cord plasma (top), non-pregnant women (middle) and pregnant women (bottom), with the percent occupancy on the y-axis and the glycan composition on the x-axis. Numbers on top indicate the glycosite.* ***B)*** *The glycosylation difference between groups per glycosite, determined by using the energy distance. Both panels show that the unique glycosylation of IgM is specific to umbilical cord plasma and is not influenced by pregnancy, most prominently on Asp46 and Asp209.*

***Supplemental Table 1: matched mother – umbilical cord plasma cohort***

| *Sample number* | *Days Pregnant* | *Comment* |
|---|---|---|
| *1* | 268 | |
| *2* | 276 | |
| *3* | 270 | |
| *4* | 260 | |
| *5* | 265 | |
| *6* | 270 | |
| *7* | 263 | |
| *8* | 266 | Twins |
| *9* | 258 | Only mother |
| *10* | 277 | |
| *11* | 270 | |
| *12* | 252 | |
| *13* | 206 | |
| *14* | 272 | |
| *15* | 281 | |
| *16* | 273 | |
| *17* | 267 | |
| *18* | 268 | Only mother |
| *19* | 266 | |
| *20* | 270 | |
| *21* | 266 | |

***Supplemental table 2: non-pregnant women cohort***

| *label* | *age* |
|---|---|
| *1* | 20.9 |
| *2* | 20.5 |
| *3* | 37 |
| *4* | 26.6 |
| *5* | 20.2 |
| *6* | 22.8 |
| *7* | 32.3 |
| *8* | 28 |
| *9* | 35 |
| *10* | 25.3 |
| *11* | 21.1 |
| *12* | 28.8 |
| *13* | 31.2 |
| *14* | 30.1 |
| *15* | 36 |
| *16* | 32.7 |
| *17* | 36.6 |
| *18* | 34.8 |
| *19* | 31.9 |
| *20* | 20.7 |
